## Supplemental figures and tables for "Allosteric activation or inhibition of PI3Kγ mediated through conformational changes in the p110γ helical domain"

Calvin K Yip

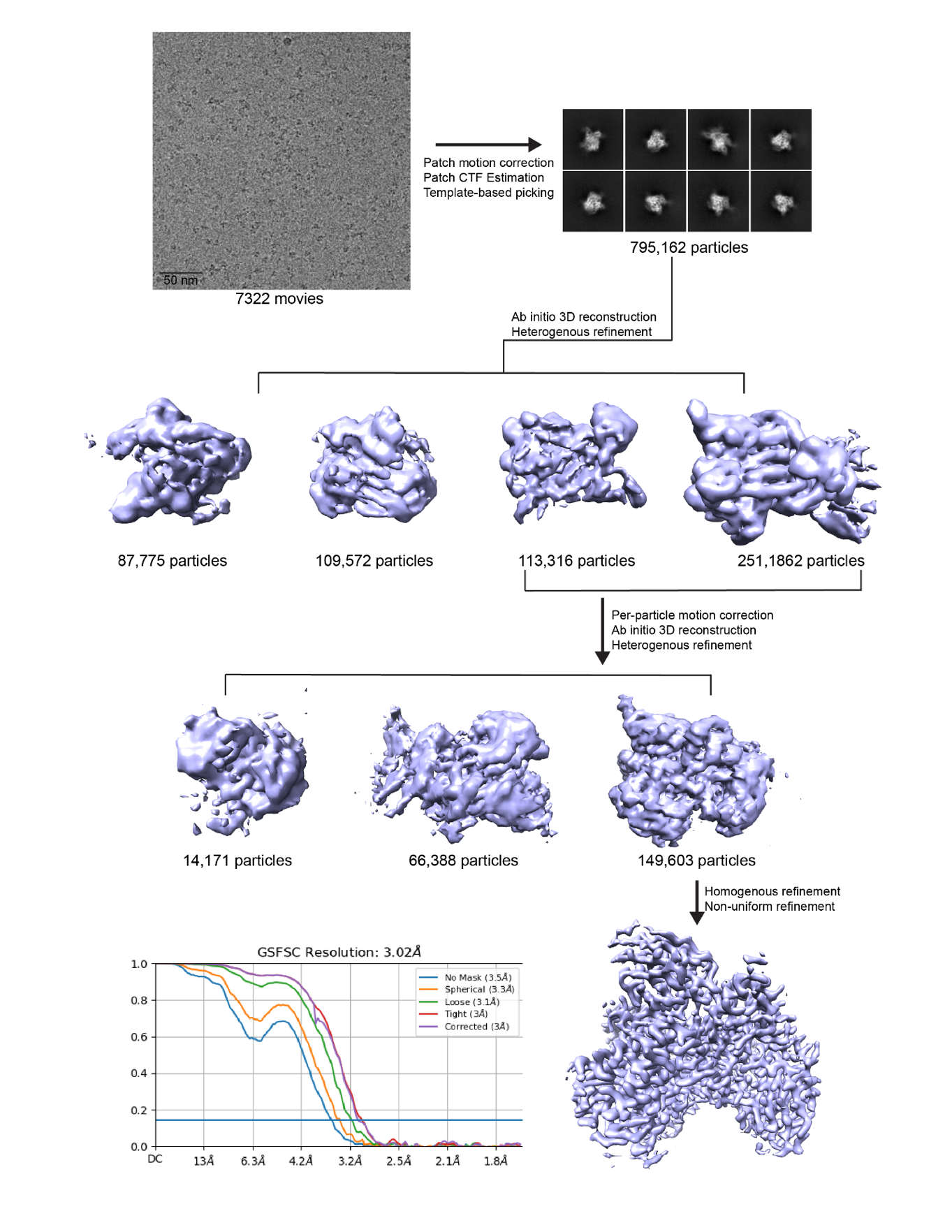

**Figure S1.** **p110**γ**-NB7 complex cryo-EM analysis workflow (related to main figure 2):** cryo-EM processing workflow of p110γ-NB7 complex are shown in order of a representative micrographs, representative 2D classification and 3D reconstruction processing strategy. Bottom left shows Gold-standard Fourier shell Correlation (FSC) curve of final round on non-uniform homogenous refinement.

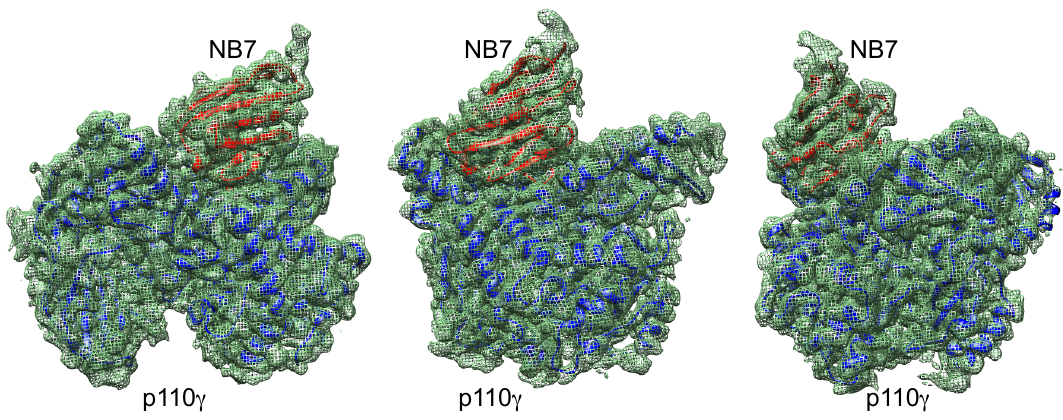

**Figure S2.** **Density fit of p110**γ**-NB7 complex (related to main figure 2):** model of p110γ (blue) / NB7 (red) complex in different orientations are shown fit within the cryo-EM density map (green mesh).

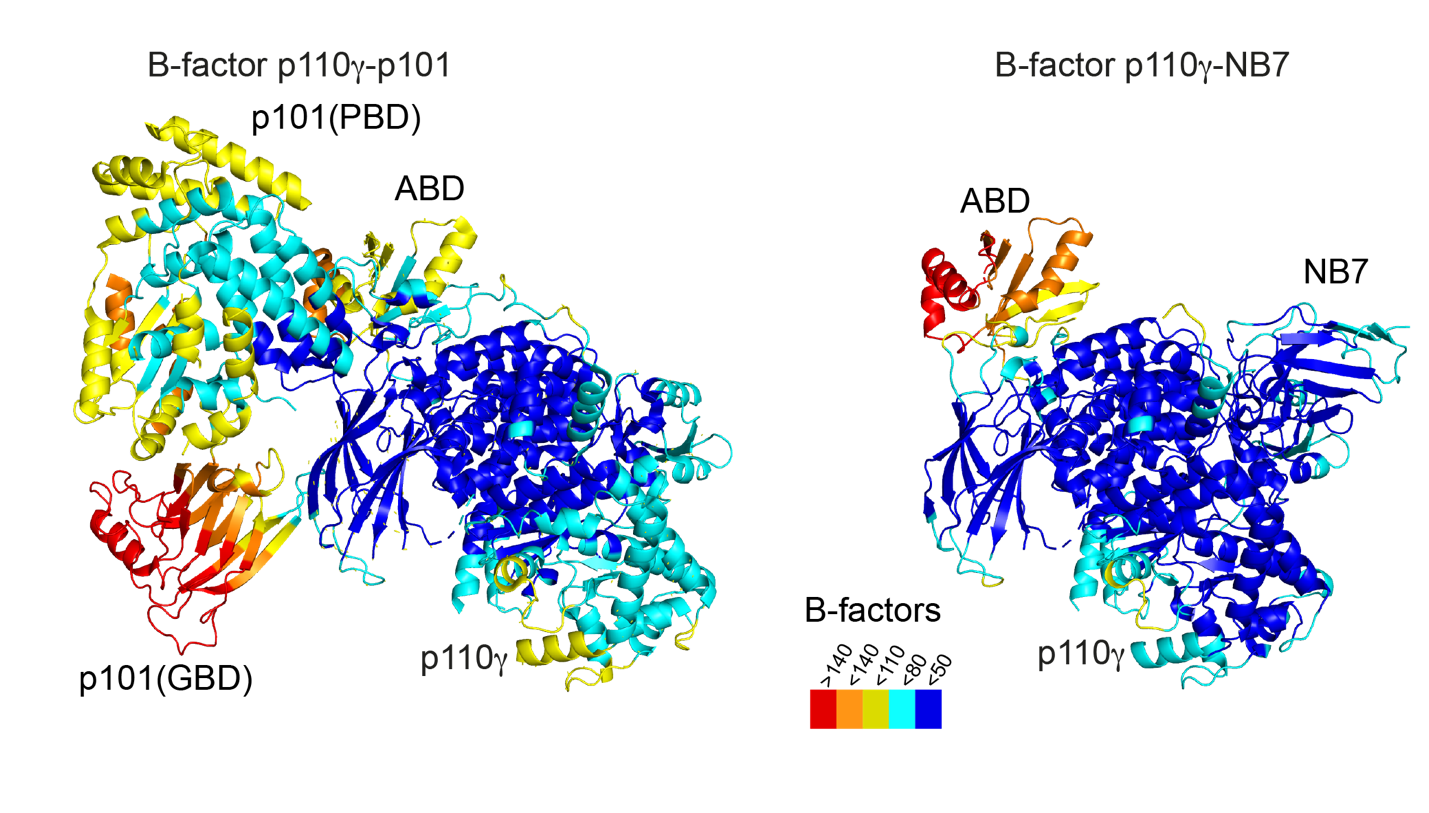
**Figure S3. Comparison of full length p110γ bound to NB7 compared to p110γ-p101 (related to main figure 2):** The structure of the p110γ-p101 complex (PDB:7MEZ) compared to the NB7-p110γ complex is shown colored according to B factor based on the legend.

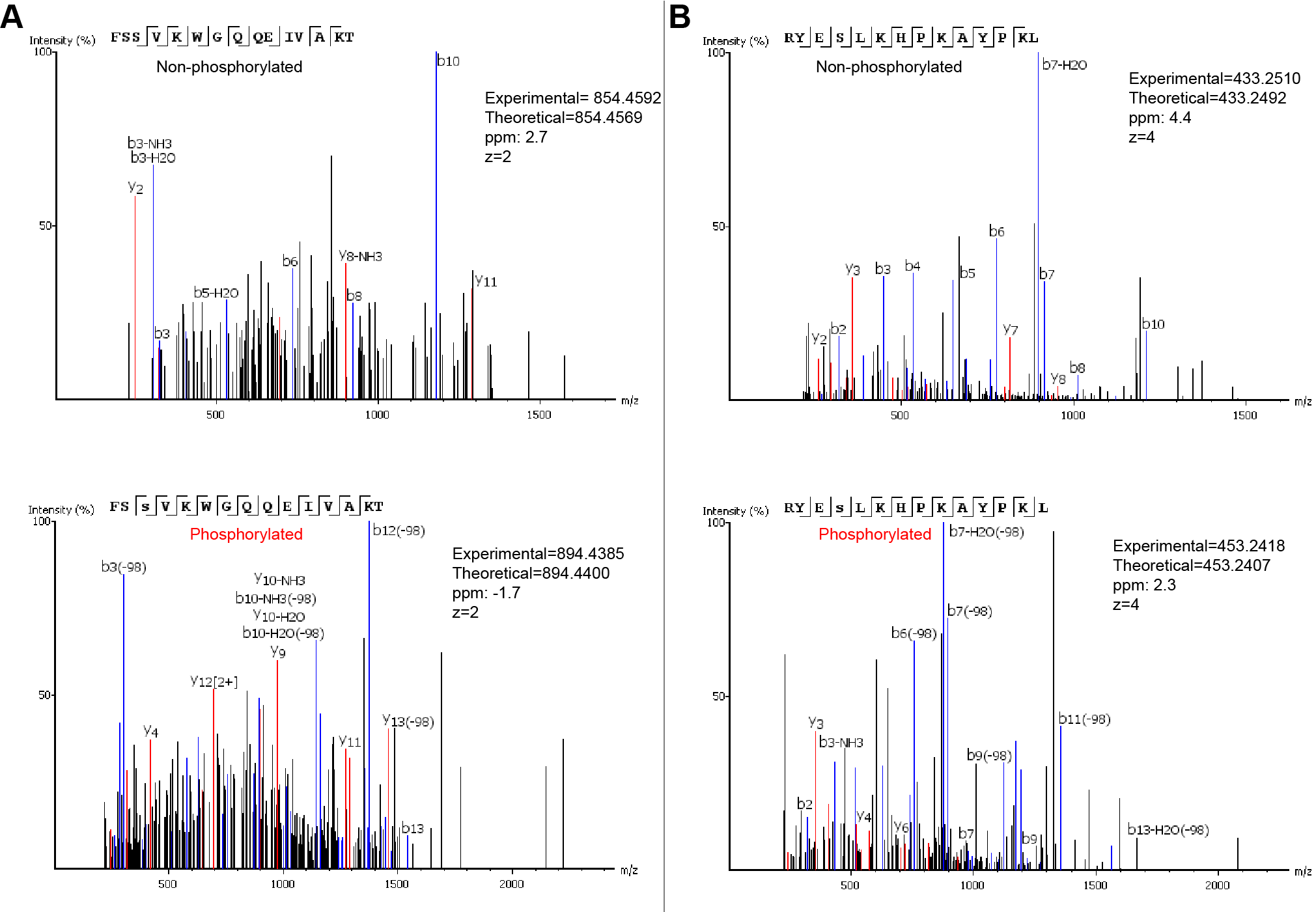

**Figure S4 (related to main figure 3).** MS/MS spectra of peptides spanning S582 and S594/S595 for both phosphorylated and unphosphorylated states. The theoretical and experimental mass are annotated for all peptides.

**Supplementary table 1.** Cryo-EM data collection, refinement and validation statistics **(related to main figure 2)**

|  | p110γ-NB7  EMD- 27627  PDB: 8DP0 |
| --- | --- |
| **Data collection and processing** |  |
| Magnification |  |
| Voltage (kV) | 300 |
| Electron exposure (e/ Å^2^) | 50 |
| Defocus range (nM) | 500-2500 |
| Pixel size (Å) |  |
| Symmetry imposed | C1 |
| Initial particle images (no.) | 795,162 |
| Final particle images (no.) | 149,603 |
| Map resolution (Å) | 3.02 |
| FSC threshold | 0.143 |
| Map resolution range (Å) | 2.6-4.4 |
| **Refinement** |  |
| Initial model used (PDB) | 7MEZ (p110γ only) |
| Model Resolution (Å) | 3.02 |
| FSC threshold | 0.5 |
| Map sharpening B factor | Sharpened locally |
| Model composition |  |
| Non-hydrogen atoms | 8737 |
| Protein residues | 1,066 |
| Ligands | 0 |
| *B*-factors |  |
| Protein | 52.4 |
| Validation |  |
| Mol probability score | 1.29 |
| Clashscore | 5.33 |
| Poor rotamers (%) | 0.0 |
| Ramachandran |  |
| Favored | 98.41 |
| Allowed | 1.59 |
| Outliers | 0.0 |
| R.m.s. deviations |  |
| Bond lengths (Å) | 0.002 |
| Bond angles (°) | 0.490 |
| Model to map fit (CC_mask) | 0.86 |

**Supplementary table 2.** HDX-MS data collection and validation statistics

**(related to main figure 4)**

| Data set | p110γ unphosphorylated | p110γ phosphorylated |
| --- | --- | --- |
| HDX reaction details | %D_2_O=75.5%  pH_(read)_=7.5  Temp=4ºC, 20ºC | %D_2_O=75.5%  pH(read)=7.5  Temp=4ºC, 20ºC |
| HDX time course (seconds) | 3s at 4ºC, 3s, 30s, 300s, 3000s at 20 ºC | 3s at 4ºC, 3s, 30s, 300s, 3000s at 20 ºC |
| HDX controls | N/A | N/A |
| Back-exchange | No correction, deuterium levels are relative | No correction, deuterium levels are relative |
| Number of peptides | 244 | 244 |
| Sequence coverage | 98.4% | 98.4% |
| Average peptide  /redundancy | Length= 15.2  Redundancy= 3.3 | Length= 15.2  Redundancy= 3.3 |
| Replicates | 3 | 3 |
| Repeatability | Average StDev=0.53% | Average StDev=0.57% |
| Significant differences in HDX | >5% and >0.4 Da and unpaired t-test ≤0.01 | >5% and >0.4 Da and unpaired t-test ≤0.01 |
